# Multiscale Modeling Uncovers Macrophage Infiltration and TNF-α Signaling Networks for Targeting in Inflammatory Breast Cancer Tumor Emboli

**DOI:** 10.1101/2025.05.29.656249

**Authors:** Pritha Pai, Christophe Van Berckelaer, Steven Van Laere, Alexandra Bennion, Theresa Charity, Jinming Yang, Francois Bertucci, Peter Van Dam, Gregory M. Palmer, Shannon McCall, Luc Y. Dirix, Naoto T. Ueno, Gayathri R. Devi

## Abstract

Inflammatory breast cancer (IBC) tumors are characterized by diffuse clusters of cells found in dermal tissue and lymphatic vessels, known as tumor emboli. Thus, IBC needs a novel treatment because it is the most aggressive breast cancer subtype. We hypothesized that the interaction between tumor emboli and the tumor immune microenvironment (TiME) fosters survival signaling, leading to the aggressiveness of the IBC. In this study, ex vivo tumor emboli were generated from patient-derived cell lines cultured in a lymphatic-like environment, which was compared to 2D monolayer cultures, revealing upregulation of TNFR signaling networks, CXCL8, and immune cell chemotaxis genes. Spatial immunophenotyping of IBC patient tumors demonstrated high levels of CD163+ tumor-associated macrophages (TAMs). Furthermore, intravital imaging of *CX3cr1^GFP^* mice confirmed macrophage movement toward tumor cell clusters. Finally, targeting macrophage-associated TNF-α-signaling using Birinapant, a SMAC mimetic, inhibited the tumor emboli phenotype *in vivo,* identifying Birinapant as a potential therapeutic agent to disrupt this signaling axis.

## Introduction

Inflammatory breast cancer (IBC) is the most aggressive form of breast cancer. IBC accounts for 10-15% of breast cancer-related deaths^1, 2^ and is typically diagnosed at stage III or higher. Notably, approximately 30% of IBC patients present with distant organ metastasis at the time of diagnosis, in contrast to about 5% in stage-matched non-IBC cases^3, 4^. Currently, there is no FDA-approved IBC-specific treatment. Patients often undergo a regimen of neoadjuvant chemotherapy, typically including taxanes or anthracyclines, followed by surgery and locoregional radiotherapy^5^. Despite advancements in trimodal therapy, the—survival rate for patients with IBC remains dismally low, under 50%^6^.

IBC does not usually present with a palpable tumor mass. Instead, a pathological hallmark is the presence of diffuse clusters of tumor cells within the breast dermal tissue and dermal lymphatic vessels, collectively referred to as tumor emboli^7–9^. These tumor clusters/emboli can obstruct lymphatic drainage, contributing to the characteristic clinical presentation of a red, swollen, and painful breast^10^.

As tumor emboli migrates through compressed lymphatic vessels, they experience both cellular and mechanical stress. This stress forces the tumor cells to adapt by upregulating survival pathways that may help them evade various stress stimuli including oxidative and immune-mediated cell death^8, 9, 11^. In this context, immunohistochemical analysis of IBC tumor emboli have shown a high expression of nuclear transcription factor NFκB, and its functional partner, the X-linked inhibitor of apoptosis protein (XIAP)^8^ both of which play key roles in survival signaling^12–14^. In addition, recent studies have shown increased infiltration of immunosuppressive cell subsets, including FOXP3+ regulatory T cells (Tregs), CD163+ tumor-associated macrophages (TAMs), and increased PD-L1 expressing T cells and the role of XIAP in fostering an immunosuppressive microenvironment^15–17^. These observations suggest that stress-adapted IBC cells with activated survival signaling pathways are involved in polarizing the tumor microenvironment toward a more immunosuppressive state. However, the precise signaling dynamics governing these interactions and how they affect IBC biology, including tumor emboli are poorly understood. Despite data generated from various molecular profiling studies in past decade, identifying the intrinsic molecular mechanisms underlying the aggressive invasion and rapid growth of IBC tumor emboli remains a challenge^18–20^. Although numerous preclinical *in vitro* and *in vivo* models have been developed to simulate tumor emboli^8, 9, 13, 21–25^, the rarity of IBC cases in a single clinical practice and the difficulty in obtaining biospecimens with clear evidence of tumor emboli limit research efforts.

The aim of this study was to conduct an in-depth interrogation of the tumor emboli and their microenvironment using transcriptomic, proteomic, and spatial immunophenotyping techniques. The primary goal was to identify opportunities for targeting this disease phenotype to inhibit metastatic progression. Furthermore, multi-scale analysis of the tumor immune microenvironment utilizing both preclinical models and clinical samples has the potential to advance our understanding of this understudied breast cancer subtype for improved clinical outcomes.

## METHODS

### Breast Cancer Samples and Expression Profiling

We queried the publicly available gene expression data from patients with IBC with documented lymphovascular invasion (N=78). These data are part of the largest series that was collected by the Inflammatory Breast Cancer International Consortium (IBC-IC) as described previously^26, 27^. Constituent data sets have been made publicly available (Antwerp: E-MTAB-1006; Marseille: E-MTAB-1547; and Houston: E-GEOD-22597). Gene expression differences between patients with and without lymphovascular invasion were calculated using generalized linear models using the BioC-package limma using only the presence of tumor emboli as an independent variable. Then, the vector of gene-wise fold-changes was used to evaluate enrichment of gene sets of interest using the BioC-package fgsea.

### Cell lines

SUM149PT and SUM190PT () cells obtained from Asterand Inc. (Detroit, MI), and MDA-IBC3 PDX kindly gifted by Dr. Wendy Woodward at MDA Anderson (Houston, TX) were cultured in Ham’s F12 medium (Mediatech Inc., Manassas, VA) supplemented with 5% FBS (Atlanta Biologicals, Lawrenceville, GA), 1% penicillin/streptomycin, 1% antibiotic/antimycotic, hydrocortisone (Invitrogen, Carlsbad, CA) and an insulin/transferrin/selenium cocktail (Gibco, Carlsbad, CA). rSUM149, a drug-resistant variant, is routinely maintained in normal culture medium supplemented with 7.5 μM research grade lapatinib (GW572016, Selleckchem, Houston, TX). The resistance reversal model rrSUM149 described previously ^28, 29^ was generated from rSUM149 after removal of lapatinib >3 months and cultured in regular SUM149 media. All cell lines are authenticated every two years and were cultured at 37°C with 5% CO_2_ for no more than 2 months prior to use in this study.

### Spatial analysis of patient samples

As part of a large effort to evaluate the immune microenvironment of IBC, we selected 176 female patients (all aged over 18 years) diagnosed with IBC enrolled in GZA Sint-Augustinus, Antwerp; Antwerp University Hospital, Antwerp and Institut Paoli-Calmettes, Marseille since 1997. All included cases had complete hospital records, histopathological confirmation of invasive carcinoma, and met the clinical diagnostic criteria for IBC as defined by international consensus^1^. Pretreatment diagnostic biopsy H&E slides from all patients were evaluated for the presence of tumor emboli and whether sufficient tissue was available for subsequent immunohistochemical analyses. Ultimately, 24 patients were included. Evaluation of the H&E samples to detect emboli was done by a board-certified pathologist specialized in breast pathology.

To analyze the composition of the immune infiltrate in the emboli of these 24 patient samples, we used the following validated antibodies: CD79α (clone: JCB117) for B-cells and plasma cells, CD8 (clone: C8/144B) for cytotoxic T-cells, FOXP3 for Tregs (clone: 236A/E7), and CD163 for TAMs (clone: MRQ-26). Staining was performed on Bond/Leica or Benchmark/Ventana auto stainers and visualized using diaminobenzidine (DAB). Slides were digitalized and evaluated using VISIOPHARM® software. With specific image analysis algorithms for every staining, we quantified the relative marker area (RMA, %) and the number of stained cells (density, #/mm²) in the emboli and tumor area. Furthermore, we also evaluated the presence of carcinoma cells immediately adjacent to the emboli.

### RNA Isolation and Sequencing

RNA from tumor cells and tumor emboli cultures were extracted as previously described previously^29^ and quantified using Qubit 2.0 Fluorometer (Life Technologies, Carlsbad, CA, USA). RNA integrity was checked using Agilent TapeStation 4200 (Agilent Technologies, Palo Alto, CA, USA). RNA sequencing libraries (n=15) were prepared using the NEBNext Ultra II RNA Library Prep Kit for Illumina following the manufacturer’s instructions (NEB, Ipswich, MA, USA). Briefly, mRNAs were enriched with Oligo(dT) beads and fragmented for 15 minutes at 94°C. First strand and second strand cDNAs were subsequently synthesized. cDNA fragments were end-repaired and adenylated at 3’ ends, and universal adapters were ligated to cDNA fragments, followed by index addition and library enrichment by limited-cycle PCR. The sequencing libraries were validated on the Agilent TapeStation (Agilent Technologies, Palo Alto, CA, USA), and quantified by using Qubit 2.0 Fluorometer (Invitrogen, Carlsbad, CA) as well as by quantitative PCR (KAPA Biosystems, Wilmington, MA, USA). Library preparation was conducted at Azenta US, Inc (South Plainfield, NJ, USA). The sequencing and analysis were conducted similar to previously described^29^.

### Gene Expression Analysis

Raw reads were mapped onto the reference genome (hg38) using the splice aware aligner HISAT2. Resulting SAM-files were converted into BAM-files and position sorted using samtools. Reads overlapping with the genomic positions of genes were counted using summarise Overlaps-function in the BioC-package GenomicAlignments. Next, genes with more than 10 counts in at least 20% of the samples were retained, vouching for a total of 14,540 unique genes.

Differences in gene expression were analyzed using negative binomial models (BioC-package DESeq2). The design matrix was set up using a model containing only the culture condition (i.e. 2D *vs*. 3D) as independent variable. Only genes with a False Discovery Rate (FDR)-corrected p-value inferior to 10% were considered differentially expressed. Results were represented in volcano plot format. List of differentially expressed genes were compared to those identified in our previous study^29^ using the Jaccard Index (JI; BioC-package GeneOverlap). In addition, Pearson correlation coefficients were used to compare vectors of log2-transformed fold-changes in this study with profiles of expression changes from previous analyses^29^ . Finally, vectors of log2-transformed fold-changes were analyzed using Gene Set Enrichment Analysis (GSEA) (BioC-package fgse) for hallmark gene sets and gene sets associated with chemotaxis of various subsets of immune cells.

### Protein Expression Analysis

Proteomics data was processed using Spectronaut v. 15.5.211111.50606 (Biognosys). Raw files were converted to .htrms using HTRMS converter. A spectral library was generated using direct-DIA searching with Spectronaut Pulsar and a SwissProt database with homo sapiens taxonomy, and with addition of porcine trypsin and BSA (20,385 entries). Database searching used default parameters, including trypsin/P specificity, up to 2 missed cleavages, N-terminal protein acetylation and Met oxidation. For protein quantification, htrms files were analyzed in Spectronaut using default settings except that workflow settings used iRT profiling of non-identified precursors, protein quantification used the MaxLFQ algorithm and q-value filtering with 0.2 percentile. Local normalization was applied using precursors identified in all runs (q-value complete).

Prior to analysis, between array quantile normalization was performed using the limma package. Differences in protein expression were analyzed using generalized linear models containing only the culture condition (i.e. 2D *vs.* 3D) as an independent variable. Only proteins with an FDR-corrected p-value <0.1 were considered differentially expressed. Results were represented in volcano plot format. List of differentially expressed proteins were compared using the Jaccard Index (JI; BioC-package GeneOverlap) and vectors of log2-transformed fold-changes using the Pearson correlation coefficient.

### Tumor emboli organoid platform

IBC cells and PDX were seeded (10K) and cultured in flat-bottom, ultra-low attachment 24-well plates (Corning, Corning, NY) containing 1ml media supplemented with 2.25% PEG-8000 (Sigma, St. Louis, MO) on a rotational platform and cultured at 37°C with 5% CO_2_ for 0-10 days. Treatments included Birinapant (APExBIO, Boston, MA) at 1000nM dose for 3 days similar to previously reported studies^17, 30^. To measure the capacity of this culture platform to simulate the characteristics of the dermal lymphatic vessel^31, 32^, viscosity measurements included kinematic viscosity using a Cannon Semi Micro Viscometer (CMSMC-25, Cannon Instrument Company, Cannon Instrument Co., State College, PA) with a calibrated range of 0.4 to 2 cSt was used. A water bath was maintained at 37.9°C by a circulating water pump (1122s, VWR, Randor, PA, US). By keeping the water bath at 37.9°C, the L beaker of water that was placed inside had a constant temperature of 37°C, calibrated to account for heat loss to the environment. The viscometer was submerged to the appropriate depth in the beaker and the temperature monitored with a second thermometer. Additionally, after adding the media to the viscometer, the system was left to acclimate to the temperature for 10 minutes. Water was used as the standard for calibration, wherein a self-calibrated constant of 0.002302 mm^2^/s^2^ was calculated to match the known kinematic viscosity of water at 37°C. The result of multiplying this calibration constant with the recorded time for each liquid was the kinematic viscosity.

In order to find the absolute (dynamic) viscosity of the media, a rearranged version of Equation AA was used:

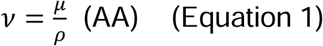

where ν is kinematic viscosity of the media (mm^2^/s or cSt), μ is absolute (dynamic) viscosity (Pa*s or cP), and ρ is density (g/mL or g/cm^3^). The solved absolute viscosity can then be utilized in later shear stress equations (Equation 2).

In order to find the densities of the media, the media were warmed up in the same method described above to 37°C. A given volume (1mL) of media (n=3) was taken and weighed using an analytical balance (Denver Instruments, Bohemia, NY, US). The mass was then divided by the volume to get density (g/mL).

In order to analytically approximate the shear stress contributed by the orbital shaker, a value for the maximum wall shear stress was calculated with Equation BB^33^:

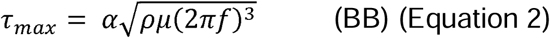

where a is the radius of rotation for the orbital shaker (2.54 cm), p is the density of the fluid (0.9899 g/mL), µ is the dynamic viscosity of the fluid (0.7124 cP = 0.007124 g/cm-s), and f is the frequency of rotation of the orbital shaker (0.667 rps = 40 rpm).

Though the orbital shaker does not apply uniform shear stress, at lower frequencies (40-70 rpm) the shear stress is relatively stable and can be treated as near-uniform^33^. Additionally, while Equation 2 is a well-cited analytical approximation for orbital-shaker setups, Ferrell, Cheng, Miao, Roy and Fissell ^33^ suggest that at lower frequencies, the equation overestimates actual shear stress by two-fold. As such, the resultant value was also halved and both results compared to target shear stress ranges.

### *In vitro* tumor emboli imaging

Tumor emboli were imaged daily for 3-6 days at 4X and 10X magnification using the light microscope EVOS M7000 (ThermoFisher, Cincinnati, OH) with phase contrast and fluorescence (GFP/RFP depending on the cell line) channel at a single plane, positioned approximately at the equator of the emboli.

### Generation of *CX3cr1^GFP^* Transgenic Mouse

Transgenic macrophage-receptor nude mice model (*Cx3cr1^GFP^* Nu/Nu) were generated wherein mice exhibit green, fluorescent macrophages. *Cx3cr1^GFP^*expressing GFP in monocytes, dendritic cells, NK cells, and brain microglia under control of the endogenous *Cx3cr1* locus were crossed with an outbred athymic mouse stocked from Jackson Laboratories (Strain #:007850) at the Duke University Breeding Core Facility (Durham, NC, USA). Athymic mice were used in this cross as they do not produce mature T cells and exhibit a reduced number of circulating lymphocytes, compared to wildtype controls. The mice were housed in a germ-free environment at the Duke Laboratory Animal Resources (DLAR). All animal studies were performed in accordance with protocols approved by the Duke University Institutional Animal Care and Use Committee (IACUC) and adhere to the NIH Guide for the Care and Use of Laboratory Animals.

### Implantation in the Window Chamber Surgical Model

Following three days in the culture platform, the tumor emboli/ organoid cultures were pooled (n=10-12), fractionated using 100 μm cell strainers (VWR, Radnor, PA), followed by trypsinization with 0.25% trypsin-EDTA for 3 minutes to prepare single cell suspensions. After neutralizing trypsin with media, the dissociated cells were washed (1X PBS) and prepared for implantation. Cells were treated with birinapant [1000nM] for 4hrs, trypsinized with 0.25% trypsin-EDTA for 2 minutes followed by neutralizing trypsin with media. Cell suspensions were centrifuged, dissolved in PBS [1X] and prepared for implantation in mice. During the imaging process, mice were anesthetized under 1-2% isoflurane (Patterson Veterinary Supply, Greeley, CO) in 100% O_2_ carrier gas for imaging. A heating pad set at 42°C kept the mice warm (internal temperature around 38°C) for the duration of each imaging session (∼30min/session). Ophthalmic ointment (LubriFresh P.M., Major Pharmaceuticals, Livonia, MI) was placed on the eyes. A custom frame – the approximate size of a microscope slide – was bolted onto the extended bolts of the window chamber so that the window remained steady while imaging. Mice were imaged at several timepoints. Images were taken on the day of surgery, ranging from 0h to 6h post-surgery. Mice were then imaged daily post-surgery for 5-10 days, after which they were humanely sacrificed.

### Intravital Imaging

Mice were imaged on an inverted microscope (Zeiss Axio Observer Z1, Carl Zeiss Microscopy, White Plains, NY) at 5X magnification (Zeiss FLUAR 5x/0.25) using a 16-bit air-cooled CMOS array (Orca Flash 4.0, Hamamatsu, 2048x2048 pixels, Japan). A mercury fluorescent light source (X-Cite 120Q, Excelitas Technology, Waltham, MA) was used for epi-illumination fluorescence measurements, and a halogen lamp was used for brightfield transmission images. For imaging the GFP-labeled cells, a filter cube (Semrock, Rochester, NY) with excitation wavelength of 470/40nm and emission bandwidth of 525/50nm was used. For imaging the RFP labeled cells, a filter cube (Semrock, Rochester, NY) with an excitation wavelength of 562/40 nm and an emission bandwidth of 624/40 nm was used. Brightfield images were obtained under a polarized filter (Chroma Technology Corp, Bellows Falls, VT) using the trans-illumination source. Each mouse received a 30-tile image across its window chamber so that the entire window chamber was imaged. Additionally, each mouse received a “snap” image of one tile at the center point of the tumor, as well as a 9-tile 3x3 image. The exposure time was kept constant during these tiled images while it changed over the course of the time through the week to avoid saturation. Because the window was not perfectly flat always, a z-stack was used to obtain images across 500mm that focused on 5 equally spaced planes. In the case of a notable area of interest, the apotome (Zeiss ApoTome.2, Carl Zeiss Microscopy) was used in small areas to obtain depth-resolved images. Zen Pro software (2012, Carl Zeiss Microscopy) was used to acquire all images. To track macrophage infiltration at the site of the tumor emboli over time, 9-tile 3x3 images were taken with a time lapse of ∼3.5 min at everyday intervals during the time course of each experiment.

### Image analysis

Following image capture using inverted microscope with Zen Pro software, the tiled and z-stack images were saved as conventional fluorescence and optical sectioning apotome image. The raw (.czi) apotome files were converted into 16bit .tif file on Fiji software (RRID:SCR_002285). The three channel images were split, and gray, green, and red colors were assigned to brightfield, GFP and RFP channel respectively. For conventional fluorescence images five Z-stacks were focused by “stack focuser” plug-in on Fiji software followed by merging the desired channels. Signal and artifacts in both GFP and RFP channels were removed, and brightness and contrast were adjusted with the auto function. Merged images were converted to 8bit .tif and .png images using Fiji software for quantification. To analyze tumor area, Apotome RFP images were analyzed in Fiji and normalized for exposure time using the “Process > Math” tool. Tumor regions were selected, minimizing signal-to-background, using the “Create Selection” tool to generate binary masks saved as regions of interest (ROIs). Tumor area within each ROI was measured on Fiji and analysis conducted using GraphPad Prism v9.3.1 (GraphPad Software, La Jolla, CA; RRID:SCR_002798; www.graphpad.com). For area fraction analysis, optical sectioning images were used, and the five Z-stacks were projected using Z-max projection function on Fiji software. To measure the macrophage signal, GFP-channel images corresponding to the most focused RFP-Z-slice (optical sectioning) were used. Within each ROI, fifteen 40×40-pixel (10,203 μm²) regions were sampled in both tumor stroma (TS) and invasive margin (IM) blinded to the intensity to minimize bias. GFP optical sectioning images were adjusted for brightness using the auto function and converted to 8-bit.tif files in Fiji software. A binary mask was generated using the Triangle thresholding algorithm^34^ to distinguish macrophage-GFP signal from the background. Predefined ROIs were overlaid, and area fraction was quantified using Fiji software. As the Triangle method is intensity-independent, exposure time was not expected to affect area fraction measurements.

### Fixation and Embedding

Harvested mouse skin tissue was rinsed in PBS to remove blood and fixed in 10% volumes of buffered formalin (NBF 10%) for 24 hours. Skin tissue was prepared by trimming individual samples for transfer into histology cassettes, preventing tissue crushing. The histology cassette clamp was closed and immersed into NBF for storage. Samples were not allowed to dry. Small tissue pieces <1-2mm were placed between sponges within the cassette, to prevent the skin from curling up. After fixing for 24 hours, cassettes were placed on Sakura Tissue-Tek VIP processor overnight and embedded in paraffin the following day.

### Immunohistochemistry

Samples from the murine studies were processed, embedded in paraffin, and sectioned serially at 4μm including a section for H&E staining. All other immunohistochemical stains were performed using the Leica Bond RX automated stainer (Leica Microsystems) using a Standard Operating Procedure and fully automated workflow (Histowiz, Inc Brooklyn, NY). The slides were dewaxed using xylene and alcohol based dewaxing solutions. Epitope retrieval was performed by heat-induced epitope retrieval (HIER) of the formalin-fixed, paraffin-embedded tissue using citrate-based pH 6 solution (Leica Microsystems, AR9961) for 20 mins at 95° C. The tissues were first incubated with peroxide block buffer (Leica Microsystems), followed by incubation for 30 min with the rabbit anti-CD3 antibody (Abcam Cat# ab16669, RRID:AB_443425) at 1:100 dilution or rabbit anti-CD45 antibody (Abcam Cat# ab10558, RRID:AB_442810) at 1:2000 dilution or rabbit anti-CD8 antibody (Cell Signaling Cat# CST98941, RRID:AB_2756376) at 1:600 dilution or with the rabbit anti-F4-80 antibody (Thermo Fisher Scientific Cat# 14-4801-82, RRID:AB_467558) at 1:200 dilution or with the rabbit anti-Ki67 antibody (Abcam, ab15580) at 1:800 dilution or with goat anti-Lyve-1 antibody (R and D Systems Cat# AF2125, RRID:AB_2297188) at 1:100 dilution for 20 mins, followed by DAB rabbit or goat secondary reagents: polymer, DAB refine and hematoxylin (Bond Polymer Refine Detection Kit, Leica Microsystems) according to the manufacturer’s protocol. The slides were dried, coverslipped (TissueTek-Prisma Coverslipper) and visualized using a Leica Aperio AT2 slide scanner (Leica Microsystems) at 40X.

### Statistical Analysis

Statistical analysis was performed using RStudio, Version 4.3.1 (RRID:SCR_000432) and plots were created with the ggplot package. To compare two categorical variables, chi-square tests or Fisher exact tests were used as appropriate. To compare continuous data between two groups or more than two groups, we used the Mann-Whitney and Kruskal-Wallis tests, respectively (GraphPad Prism, RRID:SCR_002798). To compare continuous variables, Spearman correlation (GraphPad Prism, RRID:SCR_002798) coefficients were used. Statistical significance was defined as *p*<0.05.

## RESULTS

### Tumor emboli phenotype exhibits gene Expression Changes Resembling Drug Resistance

The diffuse nature of IBC presentation in patients (**Figure 1A**) coupled with the small size of the core-biopsy makes it difficult to obtain biospecimens with well annotated emboli for molecular studies. Therefore, we optimized an *in vitro* culture platform (**Figure 1B**) for generating IBC-patient-derived tumor emboli/organoid cultures. Herein, the measured physiochemical parameters such as 1.136 cSt kinematic viscosity, 1.131 cP dynamic viscosity, 0.9957 g/mL density, and 1.16 dyne/cm² shear wall stress recapitulate the physiological conditions of the dermal lymphatic environment. Importantly, this platform supports long-term culture and live imaging of tumor emboli for up to 10 days, facilitating in-depth studies of tumor behavior and drug response (**Figure 1B, bottom panel**). We conducted multi-omics analysis, including transcriptomics and proteomics, of 3D tumor emboli cultures derived from the treatment-naïve SUM149 cell line—widely recognized as a representative IBC model^35^ and compared to 2D monolayer cultures. This analysis identified 8,441 differentially expressed genes (DEGs) (**Figure 2A, Table S1**) between 2D and 3D tumor emboli culture conditions, and 3,668 differentially expressed proteins, as illustrated in the volcano plots (**Figure 2B, Table S2**).

**Figure 1.**
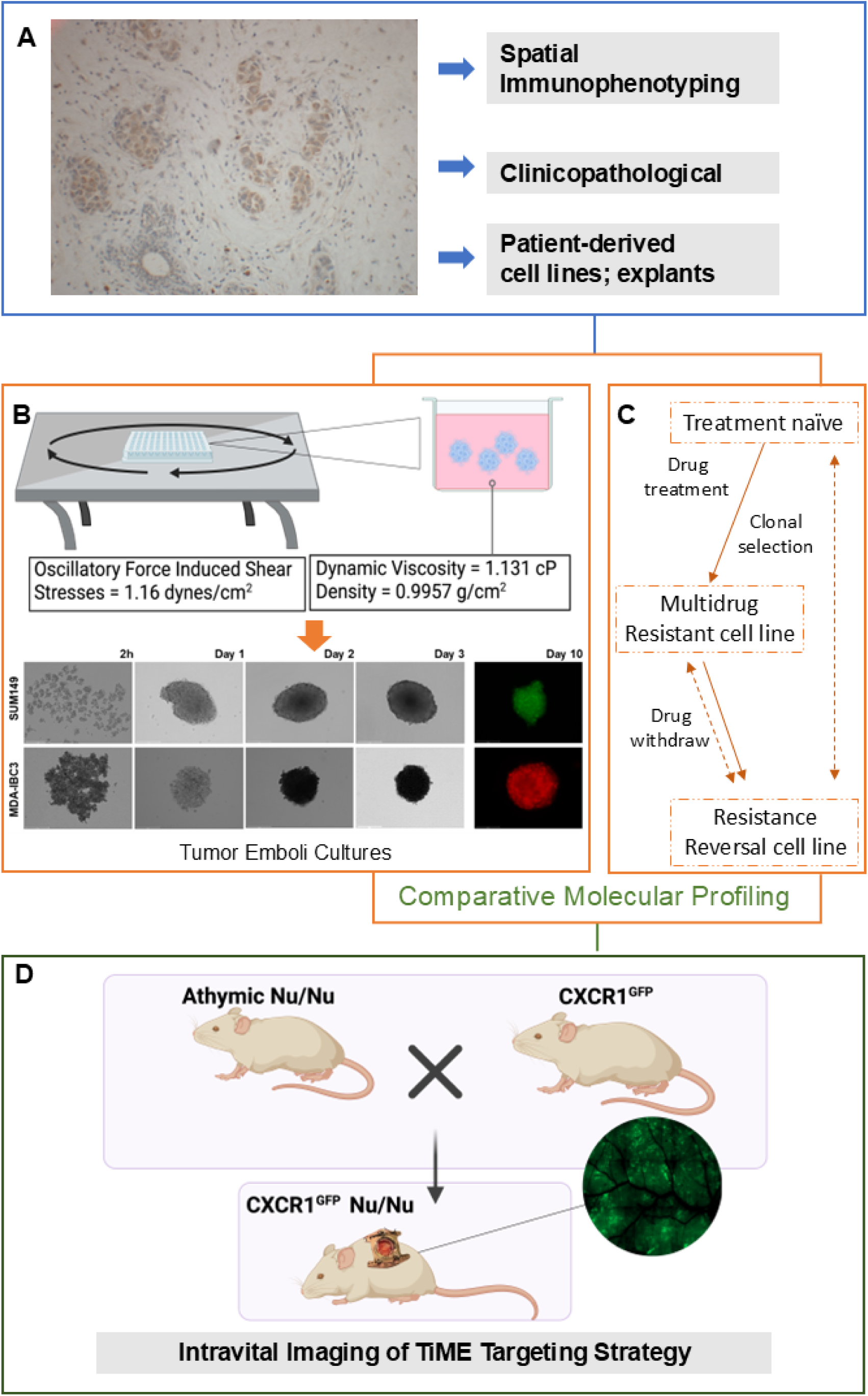
Study Models and Workflow. **A,** Representative image of IBC patient tumor emboli stained for pro-survival protein, XIAP. **B,** Tumor emboli culture platform depicting integration of biologically relevant lymphatic viscosity and shear stress measurements with representative images of IBC tumor emboli cultures maintained up to 10 days. **C,** Cell line models of acquired therapeutic resistance and reversal used for comparative analysis **D,** Intravital imaging of TiME and targeting.

**Figure 2.**
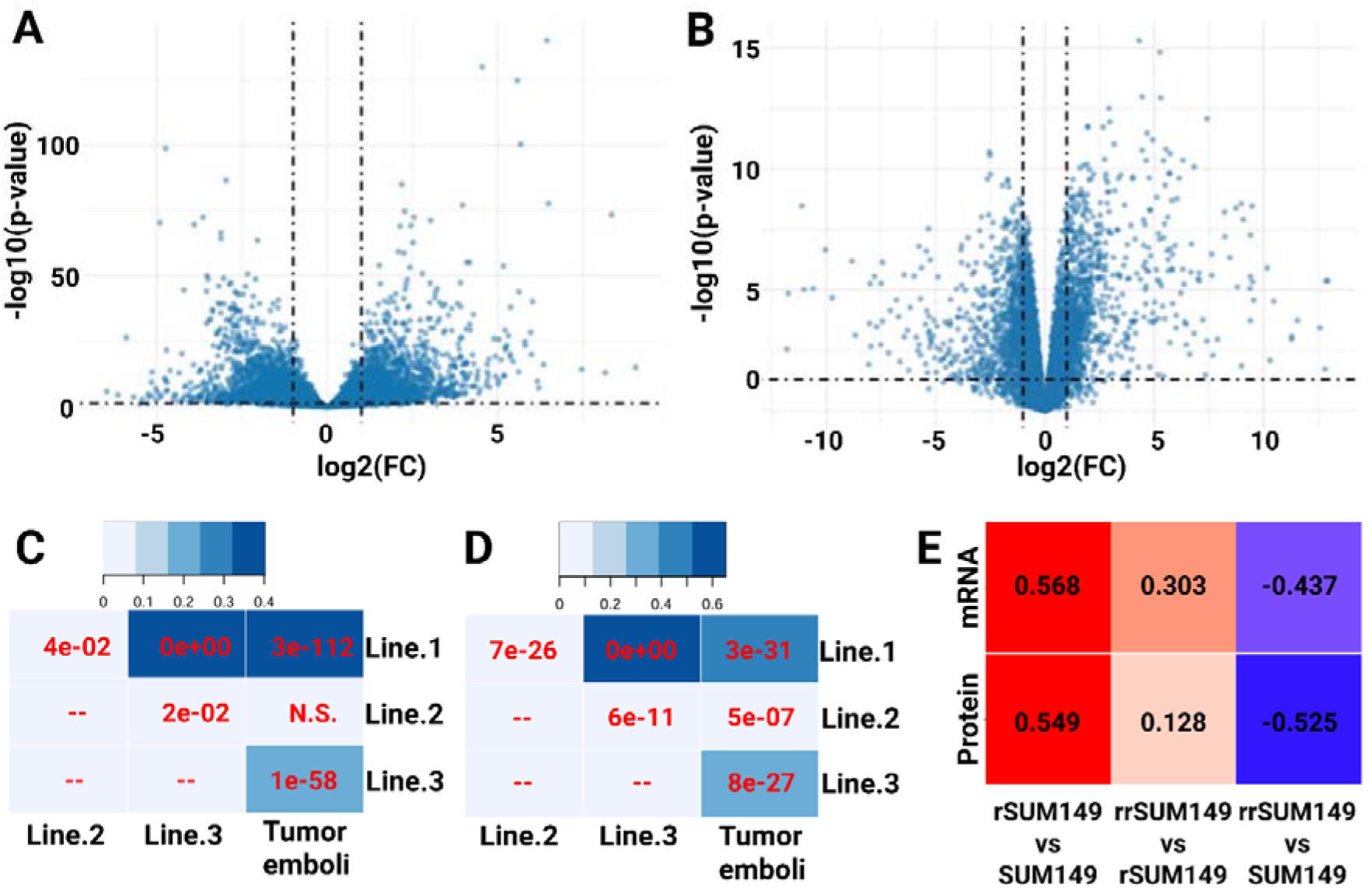
3D tumor emboli transcriptomic and proteomic analysis reveal overlapping with drug resistant IBC phenotype. **A,** Volcano plot of mRNA reveals differentially expressed genes between 2D cell lines and 3D emboli (distinct transcriptional profiles). **B,** Volcano plot of differentially expressed proteins subpopulations between 2D and 3D emboli. **C**, Comparison of gene sets using jacard index reveal overlap of transcriptomic profile between tumor emboli and drug-resistant phenotype. **D,** Comparison gene sets using jacard index reveal overlap of proteomic profile between tumor emboli and drug-resistant phenotype. **E,** Transcriptomic and proteomic analysis were conducted on SUM149 tumor emboli, SUM149 treatment naïve cell lines, resistant SUM149 cell lines (rSUM149), and resistant reversal SUM149 cell lines (rrSUM149) to calculate correlation coefficient.

Next, we compared the SUM149 3D tumor emboli cultures to a previously described^28, 29^ IBC cancer progression model that includes isogenic cell lines representing acquired multi-drug resistance to multiple FDA approved chemotherapeutics (rSUM149) as well as a cell line selected for resistance reversal (rrSUM149) **(Figure 1C).** Our analysis (**Figure 2C)** revealed that the DEGs between SUM149 and drug-resistant rSUM149 cells show substantial overlap with those differentially expressed in the 3D tumor emboli cultures (JI=0.402; P<0.001). Additionally, we observed a significant positive correlation coefficient between the vectors of log2-transformed fold-changes (R=0.568; P<0.001) (**Figure 2E**). For genes differentially expressed between rSUM149 and rrSUM149 cells (variant that has regained drug sensitivity), a less extensive albeit significant overlap was identified with genes differentially expressed between SUM149 cells cultured in 2D vs. 3D conditions (JI=0.219; P<0.001) with a negative correlation coefficient between the vectors of log2-transformed fold-changes (R=-0.437; P<0.001; **Figure 2 C, E**).

### Tumor emboli phenotype exhibits Protein Expression Changes Resembling Drug Resistance

Similarly for the proteomics data, an extensive overlap of proteins differentially expressed between SUM149 cells cultured in 2D *vs*. 3D conditions with proteins differentially expressed between SUM149 and rSUM149 cells was observed (JI=0.399; P<0.001) along with a significant positive correlation coefficient between the vectors of log2-transformed fold-changes (R=0.549; P<0.001; **Figure 2D, E**). For proteins differentially expressed between rSUM149 and rrSUM149 cells, a significant overlap was also identified with genes differentially expression between SUM149 cells cultured in 2D *vs*. 3D conditions (JI=0.337; P<0.001) with a negative correlation coefficient between the vectors of log2-transformed fold-changes (R=-0.525; P<0.001) (**Figures 2D, E**). Collectively, these results demonstrate that the switching from a monolayer to 3D tumor emboli phenotype is associated with gene expression changes that are similar to those acquired during drug resistance.

### Inflammatory and Immune signature enriched in the Tumor Emboli Cells

Gene set enrichment analysis (GSEA) on the complete vector of log2 fold changes revealed tumor necrosis factor (TNFα) signaling *via* NFκB and inflammatory responses emerging as the top two enriched hallmarks among the genes overexpressed in the 3D tumor emboli cultures **(Figure 3A)**. Further evaluation of the associated leading-edge genes revealed many chemokines and cytokines involved in chemotaxis of leukocytes such as *CCL20, IL6, CCL5, CSF1, IL1B, CSF3, CXCL2, CXCL3, CXCL11*, and *CXCL10*. Therefore, we formally tested gene ontology gene sets related to chemotaxis of leukocyte populations. Results are shown in **Figure 3B**, identifying lymphocyte (leading edge genes: *CCL20, CCL5, WNT5A, CXCL11, HSD3B7, CXCL10, NEDD9, SAA1*) and monocyte (leading edge genes: *CCL20, SLAMF8, IL6, CCL5, CXCL10, LGALS3, CREB3*) chemotaxis as the top enriched gene sets. In addition, gene sets related to chemotaxis of neutrophils, granulocytes and macrophages were significantly enriched amongst genes overexpressed in the 3D tumor emboli cultures. Collectively, these results obtained from the tumor emboli culture model are consistent with recent work from our group, revealing upregulation of TNFR1 and inflammation-related genes along with increased infiltration of CD163+ tumor associated macrophages (TAMs) in patient IBC samples with high XIAP expressing tumors^17^.

**Figure 3.**
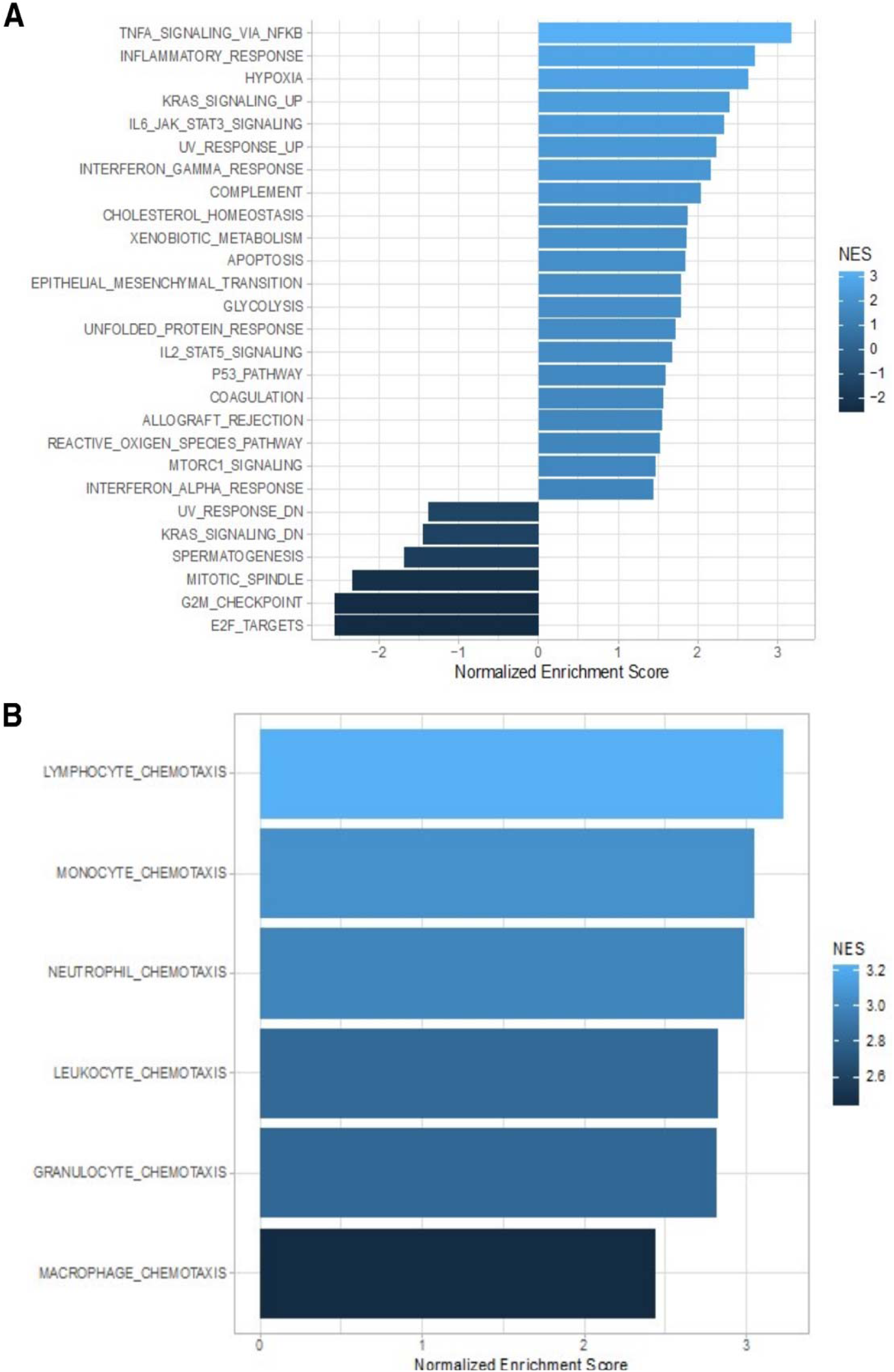
Gene set analysis of tumor emboli cultures. Highly immunogenic tumor emboli are hallmarked by (**A**) TNFα:NFκB signaling axis overexpression and (**B**) upregulation of chemotaxis of immune infiltrates, including lymphocytes, monocytes, neutrophils, leukocytes, granulocytes, and macrophages.

### Increased Infiltration of Tumor Associated Macrophages in IBC Patient Tumor Emboli

Macrophages, known for their role in TNFα production and inflammatory responses, which potentiate TNFR1-mediated survival signaling, have been identified in IBC tumors. They are also the predominant immune infiltrate in XIAP overexpressing tumors. Additionally, tumor emboli in IBC patients show significant XIAP staining (**Figure 1A**). However, no studies to date have assessed immune infiltration associated with the tumor emboli in IBC patients specifically. Because bulk expression data are not sensitive enough to detect processes in a minor cell population, we assessed the infiltration of different sets of immune cells in tumor emboli from 24 patients with IBC (**Figure 4A-B**) using immunohistochemistry for CD8 (cytotoxic T-cells), CD79α (activated B-cells), FOXP3 (regulatory T-cells) and CD163 (TAMs). We observed that TAMs are one of the most abundant immune cell types in tumor emboli present in 19 of 23 evaluated emboli (83%) biospecimens. The other cell types less commonly observed in the tumor emboli included CD8+ cells (12/24; 50%), CD79α+ cells (11/24; 46%), and FOXP3+ (6/23; 26%) cells. Furthermore, the ratio of cell densities between both compartments (emboli and stroma) was the highest for TAMs (**Table 1**).

**Figure 4.**
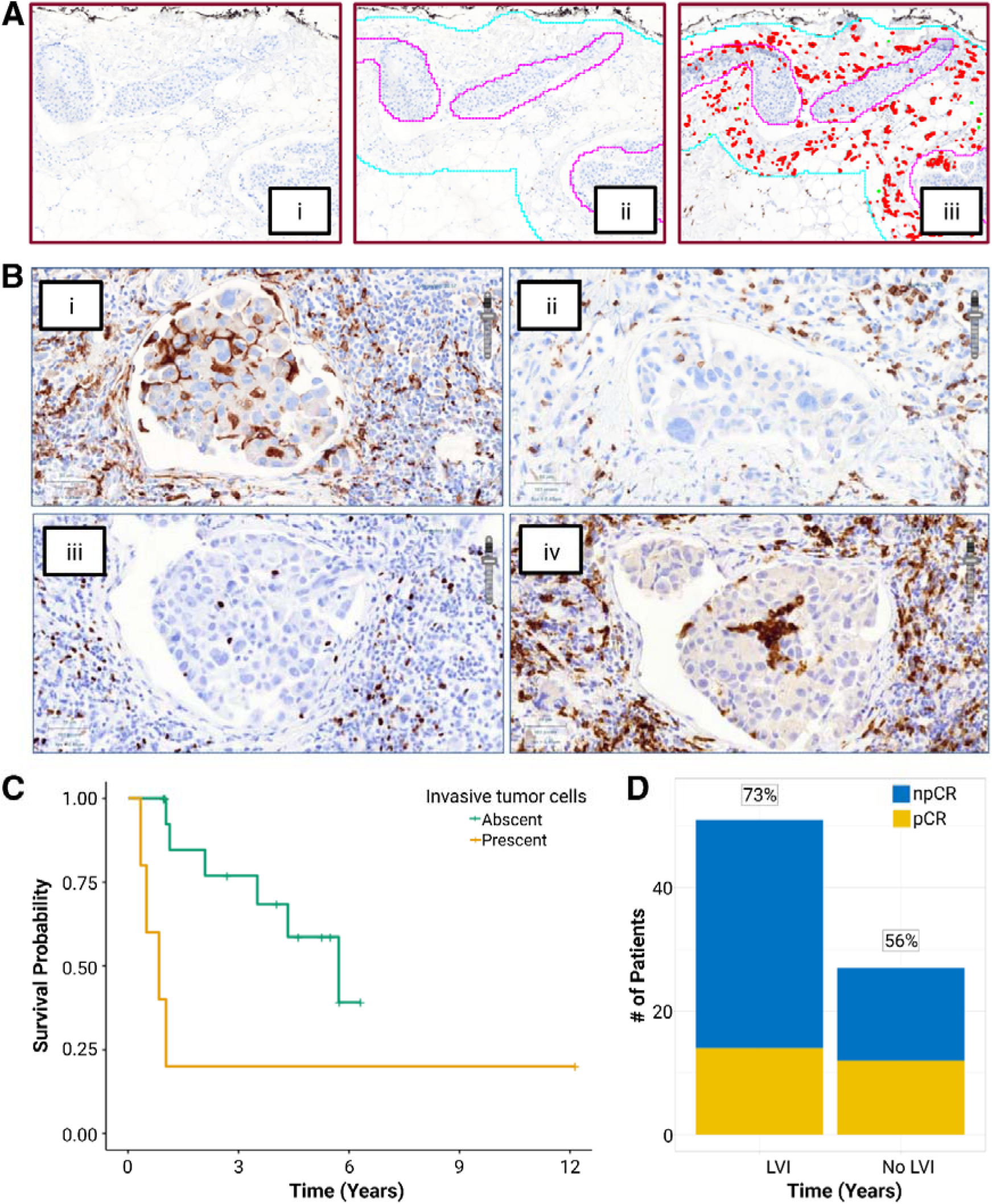
Clinicopathological and immune characteristics of IBC patients. **A,** Spatial Immunophenotyping - i. After the staining of sequentially cut slices of the marked tumor sample, slides are digitalized. ii. Tumor and emboli regions are annotated. iii Slides are analyzed using image analysis algorithms in Visiopharm®. **B,** Carcinoma cells and inflammatory cells form cohesive homotypic and heterotypic multicellular aggregates. Staining for (i) CD163, (ii) CD8, (iii) FOXP3 and (iv) CD79α. **C,** Kaplan-Meier curve demonstrating the shorter DFS (HR: 4.1, 95%CI: 1.1 – 15.0, P= 0.03) of patients with clusters of invasive tumor cells adjacent to the emboli. **D.** Bar plot showing the distribution of the number of patients (Y-axis) with (yellow) and without (blue) pathological complete response to neoadjuvant chemotherapy with respect to lymphovascular invasion (X-axis). The percentage of patients without pathological complete response per category is indicated at the top of each bar. (Odds Ratio = 2.093).

**Table 1:** Cohesive tumor emboli within lymphatic vessels are often considered to be a hallmark of IBC. We examined the immune infiltrate in these emboli in 24 IBC patients. We report the median density in the invasive carcinoma (IC), the median density in the emboli (E), the E/IC ratio and the correlation between the density in IC and E.

| | # patients | Dens (IC)<br>#/mm<sup>2</sup> | Dens (E)<br>#/mm<sup>2</sup> | Ratio Dens<br>E/IC | Correlation<br>($\rho$ & P-value) |
| --- | --- | --- | --- | --- | --- |
| **CD8** | 12/24 (50 %) | 390<br>(39 – 1663) | 37<br>(0 – 2080) | 0.18<br>(0 – 3.44) | 0.32<br>$P=0.24$ |
| **CD163** | 19/23 (83 %) | 368<br>(4 – 1085) | 196<br>(29 – 1987) | 0.51<br>(0.08 – 10.59) | 0.41<br>$P=0.10$ |
| **CD79α** | 13/24 (46 %) | 209<br>(21 – 1650) | 43<br>(6 – 2618) | 0.24<br>(0.01 – 2.98) | **0.64**<br>**$P=0.01$** |
| **FOXP3** | 6/23 (26 %) | 60<br>(3 – 314) | 9<br>(0 – 269) | 0.19<br>(0 – 9.26) | **0.62**<br>**$P=0.01$** |

Next, we characterized the ratio of densities for each immune cell type in the tumor emboli *vs*. the invasive carcinoma. Correlation analysis demonstrated that cell densities for FOXP3+, and CD79α + cells in both compartments were significantly correlated, but not for CD8+ and CD163+ cells. We confirmed these results using the RMA (**Table 2**). There was no correlation between hormone receptor status and a specific immune infiltrate in the emboli. However, a higher number of CD79α+ cells was seen in HER2+ patients (P= 0.006). Furthermore, the presence of activated B cells in the emboli also correlated with PDL1 expression (P= 0.03). Finally, more influx of sTIL in the invasive carcinoma correlated with more CD8+ (P= 0.05) FOXP3+ (P= 0.04), and CD79α+ (P= 0.01) cells in the emboli.

**Table 2:** Cohesive tumor emboli within lymphatic vessels are often considered to be a hallmark of IBC. We examined the immune infiltrate in these emboli in 24 IBC patients. We report the median RMA in the invasive carcinoma (IC), the median RMA in the emboli (E), the E/IC ratio for RMA and the correlation between the RMA in IC and E.

|  | **RMA (E) %** | **RMA (IC) %** | **Ratio RMA E/IC** | **Correlation**<br>**($\rho$ & P-value)** |
| --- | --- | --- | --- | --- |
| **CD8** | 0.08<br>(0 – 5.70) | 1.08<br>(0.06 – 5.86) | 0.17<br>(0 – 3.96) | 0.43<br>*P*= 0.10 |
| **CD163** | 0.77<br>(0.03 – 5.60) | 0.83<br>(0.01 – 3.69) | 0.74<br>(0.03 – 5.48) | 0.44<br>*P*= 0.07 |
| **CD79α** | 0.04<br>(0.01 – 10.75) | 0.57<br>(0.06 – 6.98) | 0.11<br>(0.06 – 3.80) | **0.7**<br>***P*= 0.01** |
| **FOXP3** | 0.01<br>(0 – 0.61) | 0.09<br>(0.01 – 0.61) | 0.16<br>(0 – 7.32) | **0.64**<br>***P*= 0.01** |

A complete pathological response (pCR) was seen in 8 out of 21 patients without metastatic disease at the moment of diagnosis, but there was no association with the immune micro-environment of the emboli. Next, we evaluated disease-free survival (DFS) in relation to the presence of tumor emboli in the patient samples (n=24) described above. None of the immune parameters tested were observed to be associated with DFS. However, in 5 patients (21%) we saw invasive tumor cells adjacent to the emboli and DFS in these patients was significantly shorter (**Figure 4C**) (HR: 4.1, 95%CI: 1.1 – 15.0, P= 0.03). To characterize therapeutic outcomes in IBC patients with clearly annotated tumor emboli, we identified a subset of 78 samples from the World IBC Consortium cohort. Our clinical outcome analysis in this cohort revealed that 73% of patients who exhibited tumor emboli in dermal lymphatic vessels did not achieve a pathological complete response (pCR) to neoadjuvant chemotherapy (Odds Ratio = 2.093) (**Figure 4D**). Although, this finding did not reach statistical significance (Chi-square test: P=0.145), it suggests a potential association between the presence of tumor emboli and a drug-resistant phenotype in IBC patients consistent with the molecular profile of tumor emboli cultures exhibiting a multidrug resistance phenotype.

### Targeting tumor emboli with a pan-IAP antagonist

Based on this patient biospecimen data that showed a high density of CD163+ macrophages in tumor emboli compared to the surrounding tissue, which also appeared to be independent of TAM density in the invasive carcinoma, we hypothesized that tumor-associated macrophages (TAMs) are locally recruited to the emboli in an active manner, rather than being passively included during emboli formation. To test this, we generated a genetically engineered athymic mice expressing GFP-tagged *CX3cr1* (fractalkine) (*CX3cr1^GFP^* Nu/Nu) to track movement of murine monocytes and macrophages (**Figure 1D**). Herein, tumor emboli cultures were injected within a window chamber surgically implanted on the dorsal skin of the mice to simulate the local tumor microenvironment and monitor the RFP (tumor) and GFP (monocytes) signals (**Figure 5Ai and 5Aii**). Data shows increased GFP signal in both tumor stroma (TS) (**Figure 5Av and S1**) and invasive margin (IM) (**Figure 5Avi and S1**) of the implanted tumor emboli by 72h (**Figure 5Aiii, 5Aiv and S2**). Immunohistochemistry analysis of the extracted window chamber sections demonstrated marked staining for macrophage marker F4/80 (**Figure 5Biii**) and the proliferative marker Ki67 (**Figure 5Bii**), as shown in representative images (**Figure 5B**).

**Figure 5.**
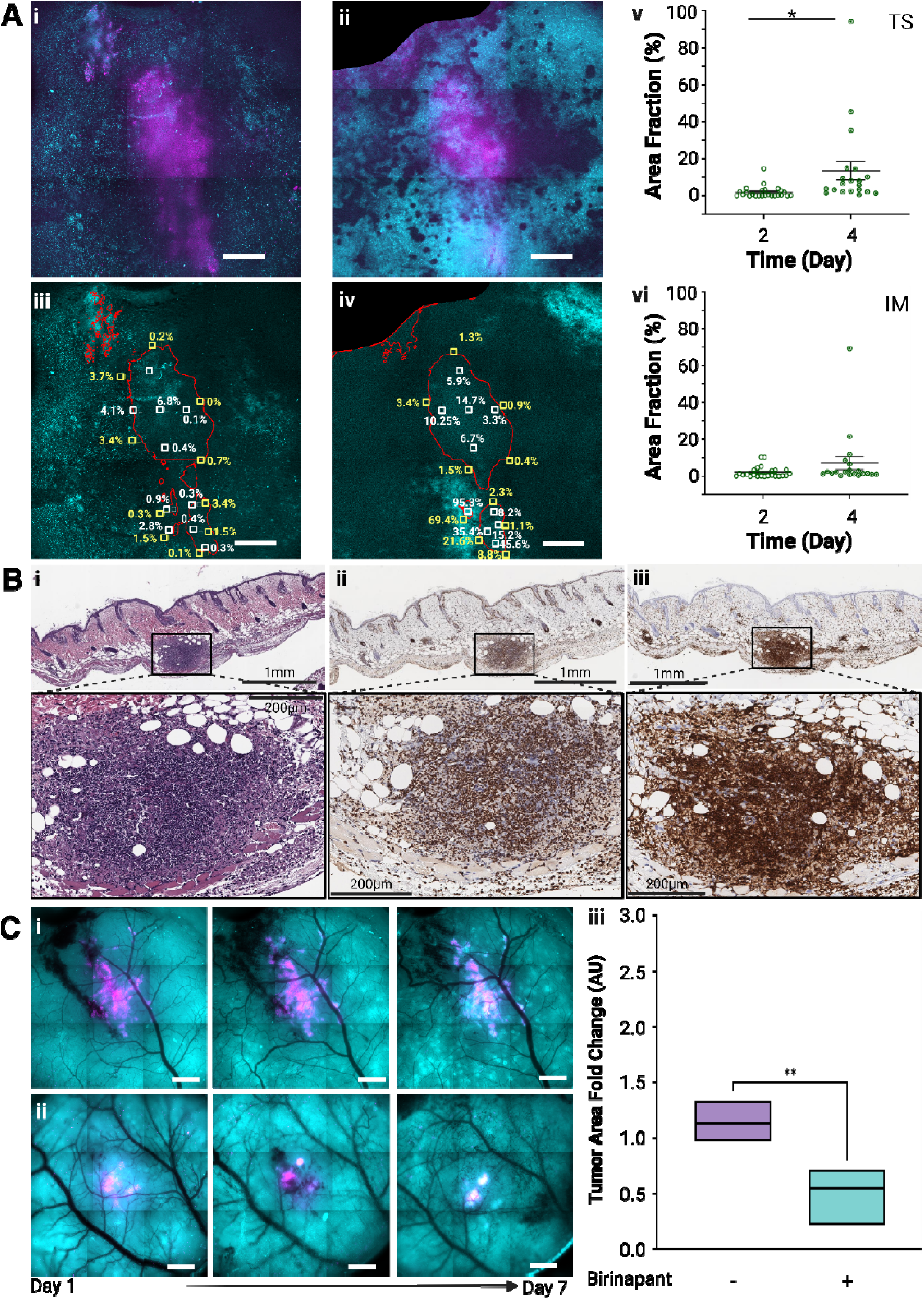
Intravital imaging. **A,** Representative Z-projection optical sectioning fluorescence images show tumor cells (RFP, red) and macrophages (GFP, green) at (i) 24h and (ii) 72h, highlighting macrophage recruitment to the tumor site. Single-plane GFP-channel images at (iii) 24h and (iv) 72h illustrate macrophage distribution. White and yellow boxes indicate tumor stroma (TS) and invasive margin (IM), respectively. **(v– vi)** Quantification of GFP+ macrophage infiltration in TS and IM regions based on area fraction above intensity threshold (each data point = 40×40 pixels, 10,203 μm²; *n* = 20; * = 0.05 > P ≥ 0.01). **B**, Immunohistochemistry of tumor emboli: (i) H&E, (ii) Ki67, (iii) F4/80 confirming IBC-like immune microenvironment. **C**, Evaluation of Birinapant treatment using window chamber model. (i) Control: SUM149-DsRed tumors with eGFP+ macrophages monitored for 7 days. (ii) Birinapant: tumors pre-treated with 1 µM Birinapant for 4h, then implanted and imaged for macrophage infiltration and colocalization. (iii) Fold change in tumor area from 0h to 144h (*n* = 8; ** =0.01>P≥0.001).**B, D**, Scale bar = 1 mm; **C inset**, scale bar = 200 µm.

Given our findings that tumor emboli attract macrophages, we sought to determine whether targeting macrophage-associated TNF-α-signaling could impair emboli-driven tumor progression. To this end, we employed the *CX3cr1^GFP^*Nu/Nu local tumor immune microenvironment model to investigate whether birinapant, a biindole-based bivalent mimetic of the naturally occurring Smac protein, previously reported^17, 36^ to inhibit TNF-α-mediated TNFR1 survival signaling could target IBC tumor emboli. Although macrophage kinetics were similar between the implanted control and Birinapant treated cells (**Figure 5C**, **S3, S4**), a significant inhibition of tumor cell spread measured by fold change in tumor area over time was observed in the Birinapant-treated group. These findings, alongside data showing enrichment of TNF-α signaling and macrophages, as well as high expression of XIAP in tumor emboli, suggest that pan-IAP antagonists like Birinapant could serve as a potential therapeutic strategy to reprogram TNF-α:TNFR1 signaling, promoting tumor cell death in IBC.

## DISCUSSION

It has been a decade since the World IBC Consortium identified a 79-gene signature for IBC, and subsequent studies, including ours, have highlighted differences in signaling pathways between IBC and non-IBC tumors and cell lines. However, our understanding of tumor emboli, the most specific characteristics of IBC, remains limited. Due to their diffuse and motile nature within dermal lymphatics, obtaining sufficient biospecimens of tumor emboli with well-annotated molecular profiles has historically been challenging. In this study, we sought to address this gap by conducting an in-depth investigation of IBC tumor emboli and their microenvironment. We employed a novel culture platform designed to simulate lymphatic shear stress, density, and viscosity to generate tumor emboli for molecular analysis, performed spatial immunophenotyping on patient samples, and developed a transgenic murine model for intravital imaging of the tumor emboli microenvironment. A key finding was the increased infiltration of macrophages within the tumor emboli along with enrichment of genes associated with TNFα signaling *via* NFκB and inflammatory responses. The efficacy of birinapant in inhibiting tumor spread in the IBC murine model further supports the potential of targeted strategies for improving immune-mediated cell death in IBC treatment.

Notably, the transition from a monolayer to a 3D tumor emboli phenotype was associated with gene expression changes that resembled those seen in acquired drug resistance. This is particularly relevant given that IBC patients with lymphatic tumor emboli, a clinicopathological marker of lymphovascular invasion, often exhibit poor responses to neoadjuvant chemotherapy. Although data was from a limited number of patient samples, it aligns with previous studies indicating that lymphovascular invasion is an independent prognostic factor in advanced breast cancers, including IBC^37, 38^.This underscores the need for multiscale models of tumor emboli to improve treatment strategies for IBC patients.

The signaling dynamics governing IBC tumor emboli and their immune interactions are still poorly understood. Unlike other cancers with lymphovascular invasion, IBC tumor emboli exhibit a distinctive geometric pattern of recurrent budding, facilitated by E-cadherin, which results in the formation of smaller daughter emboli during spreading^39^. Interestingly, macrophages activated by chemokines like IL-4 and IL-13 have been shown to promote a Th2 immunosuppressive response and express E-cadherin^40^. While it remains challenging to determine whether immunosuppressive macrophages are recruited to the tumor site or whether motile emboli translocate to immune-suppressive niches, our live imaging of macrophages and tumor cells, along with endpoint immunohistochemical analysis using the *CX3cr1^GFP^*Nu/Nu murine window chamber model, shows active recruitment of macrophages to the tumor site, with significant infiltration observed within days.

The tumor emboli culture model developed holds promise for identifying druggable pathways in IBC. A comparative analysis between this 3D culture model, treatment-naïve IBC cells, and a previously reported multi-drug resistant IBC variant^29^, revealed upregulation of genes involved in the chemotaxis of lymphocytes and monocytes. This finding supports previous reports showing that, unlike non-IBC (nIBC) tumors, which are enriched with anti-tumor immune cells such as M1 macrophages, γδ T-cells, and B memory cells, IBC tumors are characterized by an increased presence of pro-tumor macrophages (M2), as well as expanded CD4+ and CD8+ T-cell populations^16^. TAMs, the most abundant immune cells in IBC, are known to express high levels of pro-survival factors like IL-8, TNF-α, and chemokines involved in the growth-regulated oncogene (GRO) and STAT3 pathways, all of which sustain the upregulated TNFα-NFκB pro-survival signaling pathway in IBC tumor clusters and emboli^41^. Consistent with this, our studies indicate that IBC tumors, especially tumor emboli, express elevated levels of XIAP and its downstream survival partner NFkB, which drive aggressive tumor growth and invasiveness^8, 13^. We have also shown that XIAP overexpression in IBC tumors is highly dependent on non-canonical TNFα-NFκB signaling^17^, suggesting that tumor-associated macrophages may play a critical role in maintaining autocrine TNF-α signaling within tumor emboli. Beyond its role in cell survival, XIAP overexpression may also contribute to a pro-tumor immune microenvironment, promoting the persistence of tumor emboli. Elevated XIAP levels in IBC tumors have been associated with increased infiltration of immunosuppressive cell subsets, including FOXP3+ Tregs and CD163+ TAMs, as well as higher PD-L1 expression^17, 42, 43^. Moreover, lymphocyte recruitment has been shown to further enhance XIAP expression in lymphoma and leukemia models, suggesting that the immune microenvironment may actively support tumor survival^44^. These interactions highlight potential mechanisms through which tumor emboli can resist treatment and evade immune responses. Notably, IBC patient samples with high levels of tumor emboli show strong XIAP staining, suggesting that the tumor emboli secretome may recruit immunosuppressive cells *via* XIAP-mediated cross-talk between MAPK and NF-kB signaling^13^. However, we recognize that aberrant XIAP activation is likely only one of many molecular pathways contributing to tumor emboli formation and survival, these preclinical models have the potential to identify other druggable targets contributing to this immunosuppressive microenvironment.

In summary, our findings emphasize the critical role of macrophages and immune interactions in immune evasion and drug resistance in IBC (https://doi.org/10.1101/2025.05.29.656249). These insights underline the need for further research into the tumor emboli microenvironment to identify novel therapeutic targets for this understudied and so aggressive type of breast cancer. Future studies will focus on tracking monocyte and macrophage kinetics within tumor emboli to deepen our understanding of these dynamics and their potential as therapeutic targets.

## List of Abbreviations

IBC: Inflammatory breast cancer
NFkB: Nuclear factor kappa-light-chain-enhancer of activated B cells
SOC: Standard of care
pCR: Pathological complete response
LVI: Lymphovascular invasion
PDX: Patient derived xenograft
XIAP: X-linked inhibitor of apoptosis
IAP: Inhibitor of apoptosis family protein
TNFR1: Tumor necrosis factor receptor 1
TNFα: Tumor necrosis factor α
GFP: Green Fluorescence Protein
RFP: RFP Fluorescence Protein
TAM: Tumor Associated Macrophage
TiME: Tumor immuno-microenvironment
PD-L1: Programmed Cell Death Ligand 1
FOXP3: Forkhead box P3
CD163: Cluster of Differentiation 163
CD8: Cluster of Differentiation 8
CD79α: Cluster of Differentiation 79α
CX3cr1: C-X3-C motif chemokine receptor 1
DFS: Disease Free Survival
sTIL: stromal tumor infiltrating lymphocytes
RMA: relative marker area
rSUM149: Resistant SUM149
rrSUM149: Resistant Reversal SUM149
IC: Invasive Carcinoma

## DECLARATIONS

### Data Availability

The datasets related to Gene expression used in this manuscript are available from the corresponding repositories without restrictions. Gene expression data (i.e., FASTQ-files, BAM-files, and associated metadata) of the SUM149, rSUM149, rrSUM149 cell lines and 3D organoid of SUM149 have been deposited to ArrayExpress (E-MTAB-13929). Any other datasets used in this manuscript are available from the corresponding author on reasonable request.

## Acknowledgements

This work was supported in part by Department of Defense Breast Cancer Breakthrough level 2 Award W81XWH2010153 (G.R.D.), American Cancer Society Mission Boost MBG-20-141-01-MBG grant (GRD), NCI of NIH Award R01CA264529 (G.R.D.), Mitchell Scholarship (AB). We also acknowledge support for the Duke Cancer Institute shared resources/cores for Optical Molecular Imaging and Analysis (OMIA), Proteomics and Metabolomics, BioRepository and Precision Pathology Center by the P30 Cancer Center Grant NIH grant P30-CA014236. We thank Paula Newell at the Duke Surgery Substrate core service, Dr. Julie Feldstein at Histowiz and Cecile Colpaert at CORE, Antwerp, Belgium for pathological consultation, Caroline Way and Devi laboratory members for technical support and helpful discussions. Figures created with BioRender.com. Funding sources had no direct involvement in the study design; collection, analysis, and interpretation of data; the writing of this manuscript; or the decision to submit this manuscript for publication.

## Author Contributions

C.V.B., G.R.D.: conceptualization, methodology, analysis, validation, reviewing, editing, resources, writing-original draft preparation, writing— reviewing and editing, supervision. G.R.D.: funding acquisition. P.P. A.B.: methodology, analysis, validation, resources, writing—reviewing and editing. S.V.L. gene expression methodology, analysis, validation. P.P., A.B., G.P., T.C., J.Y.: murine studies, methodology, analysis; S.V.L., G.R.D., C.V.B., F.B., N.U., P.V.D., L.D., S.M.: clinical dataset, methodology, writing—review and editing. All authors have read and agreed to the published version of the manuscript.

## Conflict of interest

The authors declare no potential conflicts of interest.

## Supplementary Figures

**Supplemental Figure 1.**
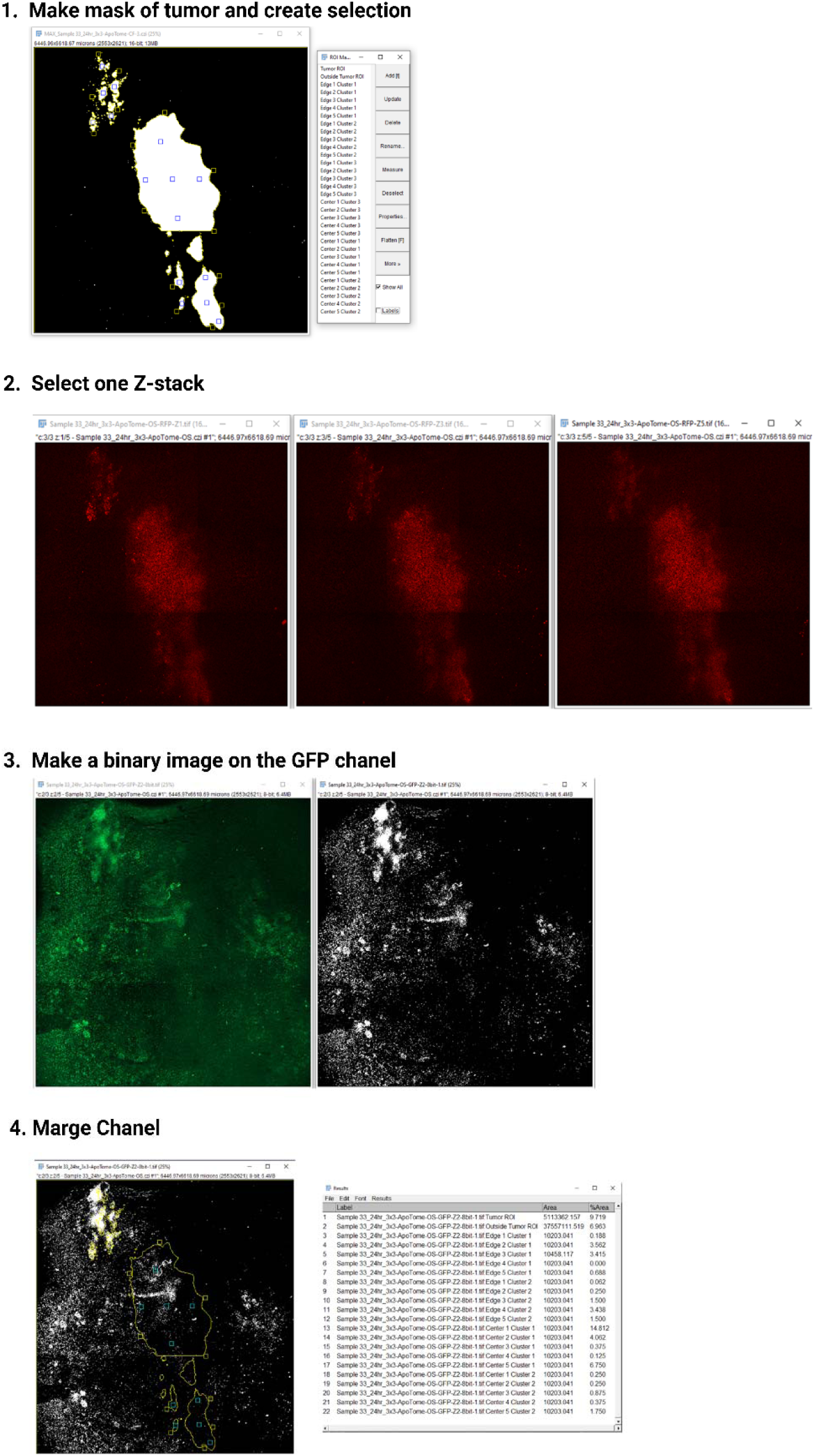
Workflow of quantitative analysis of intravital Optical Sectioning (OS) images from a representative murine window chamber.

**Supplemental Figure 2.**
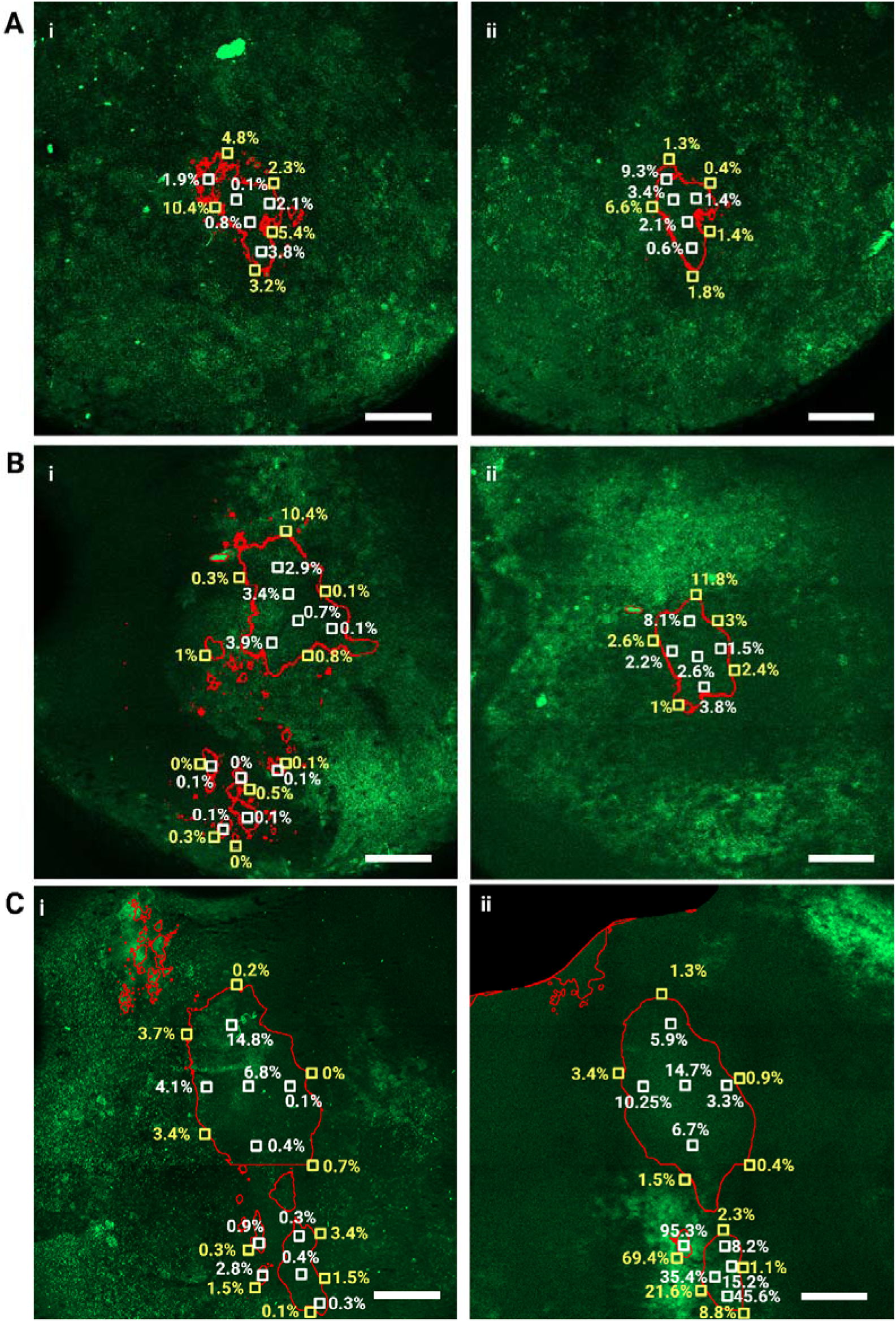
Representative single stack optical sectioning fluorescence images of GFP channel showing macrophage infiltration at (i) 24 and (ii) 72 hours with boxes [white for tumor stroma (TS) and yellow for invasive margin (IM)] area fraction measurements along with corresponding values. Scale bar represents 1mm.

**Supplemental Figure 3.**
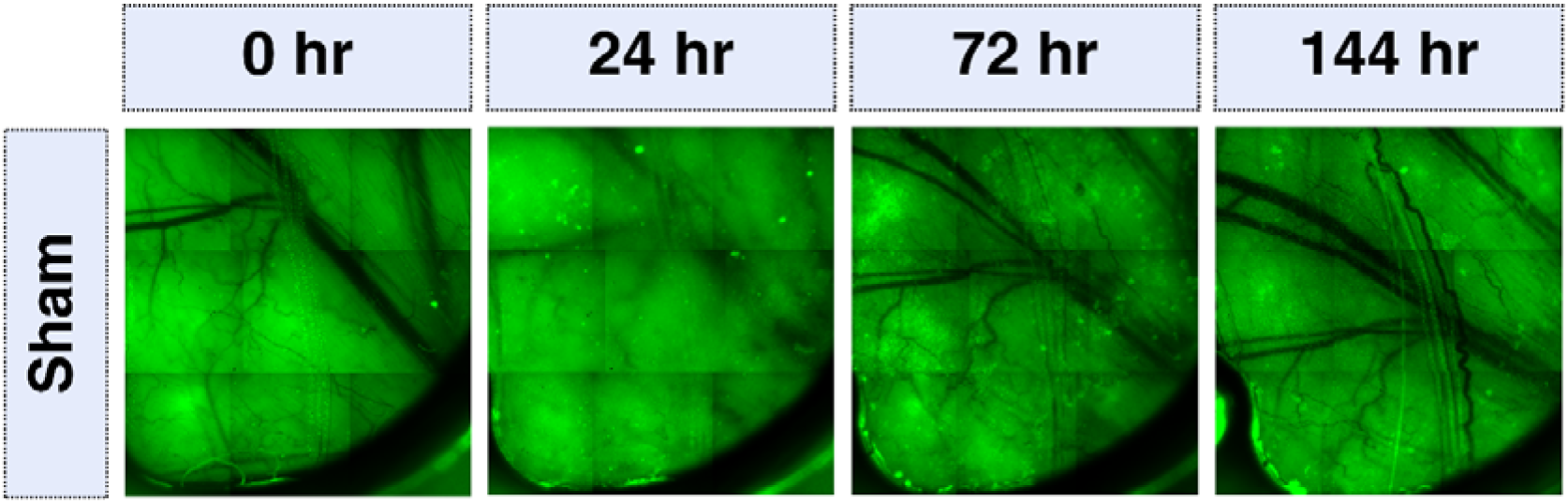
Representative photomicrographs from intravital imaging of sham (no tumor cells) window chambers from mice as controls to show lack of any macrophage signals or surgery related inflammation.

**Supplemental Figure 4.**
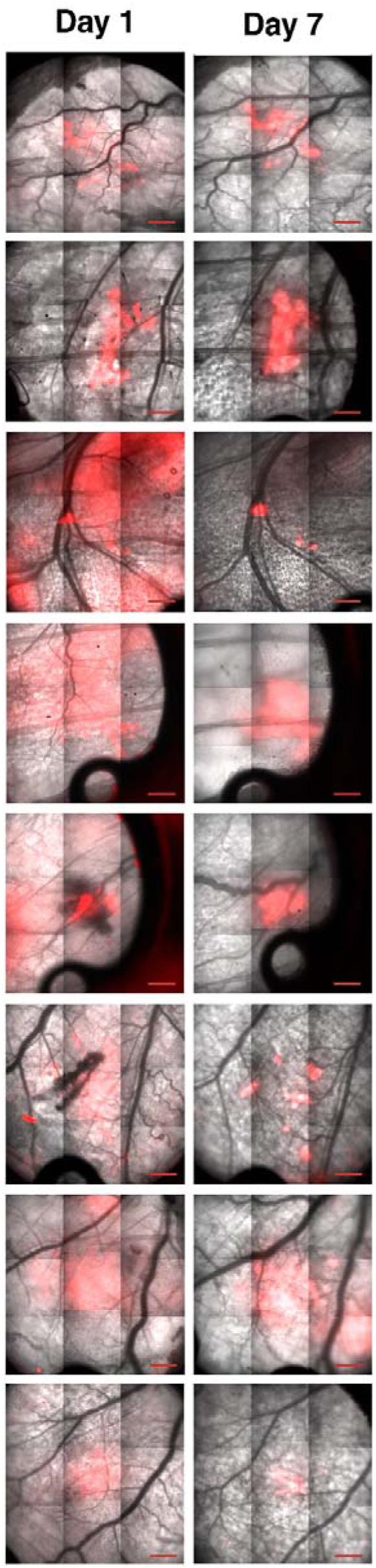
Representative intravital images (merged bright field and RFP channel images at Day 1 and Day 7) from mice window chambers implanted with birinapant treated tumor cells.

## Supplemental Tables

**Supplementary Table 1:** List of genes studied during 3D vs 2D SUM149 culture. Please see attached file “Supp. Table1”.

**Supplementary Table 2:** List of genes studied during 3D vs 2D SUM149 culture. Please see attached file “Supp. Table2”.

## Notes

### Competing Interest Statement

The authors have declared no competing interest.

### Summary of Updates

Abstract Figure 5 Legend of Table 1 Supplementary table moved as Table 2 Addition of two new supplementary tables The method section was updated by adding some details and RRIDs

## References

1. Dawood S, Ueno NT, Valero V, Woodward WA, Buchholz TA, Hortobagyi GN, Gonzalez-Angulo AM, Cristofanilli M. Differences in survival among women with stage III inflammatory and noninflammatory locally advanced breast cancer appear early: a large population-based study. Cancer. 2011;117(9):1819–26. doi: 10.1002/cncr.25682. PubMed PMID: 21509759.

2. Abraham HG, Xia Y, Mukherjee B, Merajver SD. Incidence and survival of inflammatory breast cancer between 1973 and 2015 in the SEER database. Breast Cancer Res Treat. 2021;185(1):229–38. Epub 2020/10/10. doi: 10.1007/s10549-020-05938-2. PubMed PMID: 33033965.

3. Wang Z, Chen M, Pan J, Wang X, Chen XS, Shen KW. Pattern of distant metastases in inflammatory breast cancer - A large-cohort retrospective study. J Cancer. 2020;11(2):292–300. Epub 2020/01/04. doi: 10.7150/jca.34572. PubMed PMID: 31897225; PMCID: PMC6930435.

4. Hirko KA, Regan MM, Remolano MC, Schlossman J, Harrison B, Yeh E, Jacene H, Nakhlis F, Block C, Rosenbluth JM, Garrido-Castro AC, Overmoyer BA. Dermal Lymphatic Invasion, Survival, and Time to Recurrence or Progression in Inflammatory Breast Cancer. Am J Clin Oncol. 2021;44(9):449–55. Epub 2021/06/22. doi: 10.1097/coc.0000000000000843. PubMed PMID: 34149037.

5. Newman AB, Lynce F. Tailoring Treatment for Patients with Inflammatory Breast Cancer. Curr Treat Options Oncol. 2023;24(6):580–93. Epub 20230412. doi: 10.1007/s11864-023-01077-0. PubMed PMID: 37043118.

6. van Uden DJP, van Maaren MC, Strobbe LJA, Bult P, van der Hoeven JJ, Siesling S, de Wilt JHW, Blanken-Peeters C. Metastatic behavior and overall survival according to breast cancer subtypes in stage IV inflammatory breast cancer. Breast Cancer Res. 2019;21(1):113. Epub 2019/10/19. doi: 10.1186/s13058-019-1201-5. PubMed PMID: 31623649; PMCID: PMC6798447.

7. Vermeulen PB, van Golen KL, Dirix LY. Angiogenesis, lymphangiogenesis, growth pattern, and tumor emboli in inflammatory breast cancer: a review of the current knowledge. Cancer. 2010;116(11 Suppl):2748-54. doi: 10.1002/cncr.25169. PubMed PMID: 20503405.

8. Arora J, Sauer SJ, Tarpley M, Vermeulen P, Rypens C, Van Laere S, Williams KP, Devi GR, Dewhirst MW. Inflammatory breast cancer tumor emboli express high levels of anti-apoptotic proteins: use of a quantitative high content and high-throughput 3D IBC spheroid assay to identify targeting strategies. Oncotarget. 2017;8(16):25848–63. Epub 2017/05/04. doi: 10.18632/oncotarget.15667. PubMed PMID: 28460441; PMCID: 5432221.

9. Bocci F, Gearhart-Serna, L., Boareto, M., Ribeiro, M., Ben-Jacob, E., Devi, G.R., Levine, H., Onuchic, J.N., & Jolly, M.K. Toward understanding cancer stem cell heterogeneity in the tumor microenvironment. Proceedings of the National Academy of Sciences. 2019;116(1):148–57.

10. Balema W, Liu D, Shen Y, El-Zein R, Debeb BG, Kai M, Overmoyer B, Miller KD, Le-Petross HT, Ueno NT, Woodward WA. Inflammatory breast cancer appearance at presentation is associated with overall survival. Cancer Med. 2021;10(18):6261–72. Epub 2021/07/31. doi: 10.1002/cam4.4170. PubMed PMID: 34327874; PMCID: PMC8446552.

11. Kulwatno J, Gearhart J, Gong X, Herzog N, Getzin M, Skobe M, Mills KL. Growth of tumor emboli within a vessel model reveals dependence on the magnitude of mechanical constraint. Integrative Biology. 2021;13(1):1–16. doi: 10.1093/intbio/zyaa024.

12. Aird KM, Ghanayem RB, Peplinski S, Lyerly HK, Devi GR. X-linked inhibitor of apoptosis protein inhibits apoptosis in inflammatory breast cancer cells with acquired resistance to an ErbB1/2 tyrosine kinase inhibitor. Mol Cancer Ther. 2010;9(5):1432–42. Epub 2010/04/22. doi: 1535-7163.MCT-10-0160 [pii] 10.1158/1535-7163.MCT-10-0160 [doi]. PubMed PMID: 20406946.

13. Evans MK, Brown, M.C., Geradts, J., Bao, X., Robinson, T.J., Jolly, M.K., Vermeulen, P.B., Palmer, G.M., Gromeier, M., Levine, H., & Morse, M.A. XIAP regulation by MNK links MAPK and NFκB signaling to determine an aggressive breast cancer phenotype. Cancer Research. 2018;78(7):1726–38.

14. Jost PJ, Vucic D. Regulation of Cell Death and Immunity by XIAP. Cold Spring Harb Perspect Biol. 2020;12(8). Epub 2019/12/18. doi: 10.1101/cshperspect.a036426. PubMed PMID: 31843992; PMCID: PMC7397824.

15. Lim B, Woodward WA, Wang X, Reuben JM, Ueno NT. Inflammatory breast cancer biology: the tumour microenvironment is key. Nature Reviews Cancer. 2018;18(8):485–99. doi: 10.1038/s41568-018-0010-y.

16. Bertucci F, Boudin L, Finetti P, Van Berckelaer C, Van Dam P, Dirix L, Viens P, Gonçalves A, Ueno NT, Van Laere S, Birnbaum D, Mamessier E. Immune landscape of inflammatory breast cancer suggests vulnerability to immune checkpoint inhibitors. Oncoimmunology. 2021;10(1):1929724. Epub 2021/06/10. doi: 10.1080/2162402x.2021.1929724. PubMed PMID: 34104544; PMCID: PMC8158040.

17. Van Berckelaer C, Van Laere S, Lee S, Morse MA, Geradts J, Dirix L, Kockx M, Bertucci F, Van Dam P, Devi GR. XIAP overexpressing inflammatory breast cancer patients have high infiltration of immunosuppressive subsets and increased TNFR1 signaling targetable with Birinapant. Transl Oncol. 2024;43:101907. Epub 20240227. doi: 10.1016/j.tranon.2024.101907. PubMed PMID: 38412664; PMCID: PMC10907867.

18. Devi GR, Hough H, Barrett N, Cristofanilli M, Overmoyer B, Spector N, Ueno NT, Woodward W, Kirkpatrick J, Vincent B, Williams KP, Finley C, Duff B, Worthy V, McCall S, Hollister BA, Palmer G, Force J, Westbrook K, Fayanju O, Suneja G, Dent SF, Hwang ES, Patierno SR, Marcom PK. Perspectives on Inflammatory Breast Cancer (IBC) Research, Clinical Management and Community Engagement from the Duke IBC Consortium. J Cancer. 2019;10(15):3344–51. Epub 2019/07/12. doi: 10.7150/jca.31176. PubMed PMID: 31293637; PMCID: PMC6603420.

19. Chakraborty P, George JT, Woodward WA, Levine H, Jolly MK. Gene expression profiles of inflammatory breast cancer reveal high heterogeneity across the epithelial-hybrid-mesenchymal spectrum. Transl Oncol. 2021;14(4):101026. Epub 2021/02/04. doi: 10.1016/j.tranon.2021.101026. PubMed PMID: 33535154; PMCID: PMC7851345.

20. Li X, Kumar S, Harmanci A, Li S, Kitchen RR, Zhang Y, Wali VB, Reddy SM, Woodward WA, Reuben JM, Rozowsky J, Hatzis C, Ueno NT, Krishnamurthy S, Pusztai L, Gerstein M. Whole-genome sequencing of phenotypically distinct inflammatory breast cancers reveals similar genomic alterations to non-inflammatory breast cancers. Genome Med. 2021;13(1):70. Epub 2021/04/28. doi: 10.1186/s13073-021-00879-x. PubMed PMID: 33902690; PMCID: PMC8077918.

21. Alpaugh ML, Tomlinson JS, Shao ZM, Barsky SH. A novel human xenograft model of inflammatory breast cancer. Cancer Res. 1999;59(20):5079–84. Epub 1999/10/28. PubMed PMID: 10537277.

22. Lacerda L, Debeb BG, Smith D, Larson R, Solley T, Xu W, Krishnamurthy S, Gong Y, Levy LB, Buchholz T, Ueno NT, Klopp A, Woodward WA. Mesenchymal stem cells mediate the clinical phenotype of inflammatory breast cancer in a preclinical model. Breast Cancer Res. 2015;17:42. Epub 2015/04/19. doi: 10.1186/s13058-015-0549-4. PubMed PMID: 25887413; PMCID: PMC4389342.

23. Nath S, Devi GR. Three-dimensional culture systems in cancer research: Focus on tumor spheroid model. Pharmacology & therapeutics. 2016;163:94–108. Epub 2016/04/12. doi: 10.1016/j.pharmthera.2016.03.013. PubMed PMID: 27063403; PMCID: 4961208.

24. Oladapo HO, Tarpley M, Sauer SJ, Addo KA, Ingram SM, Strepay D, Ehe BK, Chdid L, Trinkler M, Roques JR, Darr DB, Fleming JM, Devi GR, Williams KP. Pharmacological targeting of GLI1 inhibits proliferation, tumor emboli formation and in vivo tumor growth of inflammatory breast cancer cells. Cancer Lett. 2017;411:136–49. doi: 10.1016/j.canlet.2017.09.033. PubMed PMID: 28965853.

25. Rickard AG, Sannareddy DS, Bennion A, Patel P, Sauer SJ, Rouse DC, Bouchal S, Liu H, Dewhirst MW, Palmer GM, Devi GR. A Novel Preclinical Murine Model to Monitor Inflammatory Breast Cancer Tumor Growth and Lymphovascular Invasion. Cancers (Basel). 2023;15(8). Epub 20230412. doi: 10.3390/cancers15082261. PubMed PMID: 37190189; PMCID: PMC10137020.

26. Van Laere SJ, Ueno NT, Finetti P, Vermeulen P, Lucci A, Robertson FM, Marsan M, Iwamoto T, Krishnamurthy S, Masuda H, van Dam P, Woodward WA, Viens P, Cristofanilli M, Birnbaum D, Dirix L, Reuben JM, Bertucci F. Uncovering the molecular secrets of inflammatory breast cancer biology: an integrated analysis of three distinct affymetrix gene expression datasets. Clin Cancer Res. 2013;19(17):4685–96. Epub 2013/02/12. doi: 10.1158/1078-0432.Ccr-12-2549. PubMed PMID: 23396049.

27. Bertucci F, Ueno NT, Finetti P, Vermeulen P, Lucci A, Robertson FM, Marsan M, Iwamoto T, Krishnamurthy S, Masuda H, Van Dam P, Woodward WA, Cristofanilli M, Reuben JM, Dirix L, Viens P, Symmans WF, Birnbaum D, Van Laere SJ. Gene expression profiles of inflammatory breast cancer: correlation with response to neoadjuvant chemotherapy and metastasis-free survival. Ann Oncol. 2014;25(2):358–65. Epub 2013/12/05. doi: 10.1093/annonc/mdt496. PubMed PMID: 24299959; PMCID: PMC3905779.

28. Williams KP, Allensworth JL, Ingram SM, Smith GR, Aldrich AJ, Sexton JZ, Devi GR. Quantitative high-throughput efficacy profiling of approved oncology drugs in inflammatory breast cancer models of acquired drug resistance and re-sensitization. Cancer Lett. 2013;337(1):77–89. doi: 10.1016/j.canlet.2013.05.017. PubMed PMID: 23689139; PMCID: PMC3763836.

29. Devi GR, Pai P, Lee S, Foster MW, Sannareddy DS, Bertucci F, Ueno N, Van Laere S. Altered ribosomal profile in acquired resistance and reversal associates with pathological response to chemotherapy in inflammatory breast cancer. NPJ Breast Cancer. 2024;10(1):65. Epub 20240729. doi: 10.1038/s41523-024-00664-0. PubMed PMID: 39075068; PMCID: PMC11286775.

30. Allensworth JL, Sauer SJ, Lyerly HK, Morse MA, Devi GR. Smac mimetic Birinapant induces apoptosis and enhances TRAIL potency in inflammatory breast cancer cells in an IAP-dependent and TNF-alpha-independent mechanism. Breast Cancer Res Treat. 2013;137(2):359–71. Epub 2012/12/12. doi: 10.1007/s10549-012-2352-6. PubMed PMID: 23225169.

31. Breslin JW, Yang Y, Scallan JP, Sweat RS, Adderley SP, Murfee WL. Lymphatic Vessel Network Structure and Physiology. Compr Physiol. 2018;9(1):207–99. Epub 20181213. doi: 10.1002/cphy.c180015. PubMed PMID: 30549020; PMCID: PMC6459625.

32. Margaris KN, Black RA. Modelling the lymphatic system: challenges and opportunities. J R Soc Interface. 2012;9(69):601–12. Epub 20120111. doi: 10.1098/rsif.2011.0751. PubMed PMID: 22237677; PMCID: PMC3284143.

33. Ferrell N, Cheng J, Miao S, Roy S, Fissell WH. Orbital Shear Stress Regulates Differentiation and Barrier Function of Primary Renal Tubular Epithelial Cells. Asaio j. 2018;64(6):766–72. doi: 10.1097/mat.0000000000000723. PubMed PMID: 29240625; PMCID: PMC5995603.

34. Zack GW, Rogers WE, Latt SA. Automatic measurement of sister chromatid exchange frequency. J Histochem Cytochem. 1977;25(7):741–53. doi: 10.1177/25.7.70454. PubMed PMID: 70454.

35. Rypens C, Bertucci F, Finetti P, Robertson F, Fernandez SV, Ueno N, Woodward WA, Van Golen K, Vermeulen P, Dirix L, Viens P, Birnbaum D, Devi GR, Cristofanilli M, Van Laere S. Comparative transcriptional analyses of preclinical models and patient samples reveal MYC and RELA driven expression patterns that define the molecular landscape of IBC. NPJ Breast Cancer. 2022;8(1):12. Epub 2022/01/20. doi: 10.1038/s41523-021-00379-6. PubMed PMID: 35042871.

36. Lalaoui N, Merino D, Giner G, Vaillant F, Chau D, Liu L, Kratina T, Pal B, Whittle JR, Etemadi N, Berthelet J, Gräsel J, Hall C, Ritchie ME, Ernst M, Smyth GK, Vaux DL, Visvader JE, Lindeman GJ, Silke J. Targeting triple-negative breast cancers with the Smac-mimetic birinapant. Cell Death Differ. 2020;27(10):2768–80. Epub 2020/04/29. doi: 10.1038/s41418-020-0541-0. PubMed PMID: 32341449; PMCID: PMC7492458 2016.

37. Rakha EA, Martin S, Lee AH, Morgan D, Pharoah PD, Hodi Z, Macmillan D, Ellis IO. The prognostic significance of lymphovascular invasion in invasive breast carcinoma. Cancer. 2012;118(15):3670–80. Epub 2011/12/20. doi: 10.1002/cncr.26711. PubMed PMID: 22180017.

38. Kariri YA, Aleskandarany MA, Joseph C, Kurozumi S, Mohammed OJ, Toss MS, Green AR, Rakha EA. Molecular Complexity of Lymphovascular Invasion: The Role of Cell Migration in Breast Cancer as a Prototype. Pathobiology. 2020;87(4):218–31. Epub 20200709. doi: 10.1159/000508337. PubMed PMID: 32645698.

39. Modi AP, Nguyen JPT, Wang J, Ahn JS, Libling WA, Klein JM, Mazumder P, Barsky SH. Geometric tumor embolic budding characterizes inflammatory breast cancer. Breast Cancer Res Treat. 2023;197(3):461–78. Epub 20221206. doi: 10.1007/s10549-022-06819-6. PubMed PMID: 36473978; PMCID: PMC9734724.

40. Van den Bossche J, Laoui D, Naessens T, Smits HH, Hokke CH, Stijlemans B, Grooten J, De Baetselier P, Van Ginderachter JA. E-cadherin expression in macrophages dampens their inflammatory responsiveness in vitro, but does not modulate M2-regulated pathologies in vivo. Scientific Reports. 2015;5(1):12599. doi: 10.1038/srep12599.

41. Huang X, Cao J, Zu X. Tumor-associated macrophages: An important player in breast cancer progression. Thorac Cancer. 2022;13(3):269–76. Epub 20211215. doi: 10.1111/1759-7714.14268. PubMed PMID: 34914196; PMCID: PMC8807249.

42. Reddy JP, Atkinson RL, Larson R, Burks JK, Smith D, Debeb BG, Ruffell B, Creighton CJ, Bambhroliya A, Reuben JM, Van Laere SJ, Krishnamurthy S, Symmans WF, Brewster AM, Woodward WA. Mammary stem cell and macrophage markers are enriched in normal tissue adjacent to inflammatory breast cancer. Breast Cancer Res Treat. 2018;171(2):283–93. Epub 2018/06/03. doi: 10.1007/s10549-018-4835-6. PubMed PMID: 29858753.

43. Cohen EN, Gao H, Anfossi S, Mego M, Reddy NG, Debeb B, Giordano A, Tin S, Wu Q, Garza RJ, Cristofanilli M, Mani SA, Croix DA, Ueno NT, Woodward WA, Luthra R, Krishnamurthy S, Reuben JM. Inflammation Mediated Metastasis: Immune Induced Epithelial-To-Mesenchymal Transition in Inflammatory Breast Cancer Cells. PLOS ONE. 2015;10(7):e0132710. doi: 10.1371/journal.pone.0132710.

44. Vashisht M, Ge H, John J, McKelvey HA, Chen J, Chen Z, Wang JH. TRAF2/3 deficient B cells resist DNA damage-induced apoptosis via NF-κB2/XIAP/cIAP2 axis and IAP antagonist sensitizes mutant lymphomas to chemotherapeutic drugs. Cell Death & Disease. 2023;14(9):599. doi: 10.1038/s41419-023-06122-2.

